# K-fiber bundles in the mitotic spindle are mechanically reinforced by Kif15

**DOI:** 10.1101/2020.05.19.104661

**Authors:** Marcus A Begley, April L Solon, Elizabeth Mae Davis, Michael Grant Sherrill, Ryoma Ohi, Mary Williard Elting

## Abstract

The mitotic spindle, a self-constructed microtubule-based machine, segregates chromosomes into two eventual daughter nuclei. In mammalian cells, microtubule bundles called kinetochore-fibers (k-fibers) anchor chromosomes within the spindle. Chromosome segregation thus depends on the mechanical integrity of k-fibers. Here, we investigate the physical and molecular basis of k-fiber bundle cohesion. We sever k-fibers using laser ablation, thereby detaching them from poles and testing the contribution of pole-localized force generation to k-fiber cohesion. We then measure the physical response of the remaining kinetochore-bound segments of the k-fibers. We observe that microtubules within ablated k-fibers often, but not always, splay apart from their minus-ends. Furthermore, we find that minus-end clustering forces induced in response to ablation seem at least partially responsible for k-fiber splaying. We also investigate the role of the putative k-fiber-binding kinesin-12 Kif15. We find that pharmacological inhibition of Kif15 microtubule binding reduces k-fiber mechanical integrity. In contrast, inhibition of its motor activity but not its microtubule binding does not greatly affect splaying. Altogether, the data suggest that forces holding k-fibers together are of similar magnitude to other spindle forces, and that Kif15, acting as a microtubule crosslinker, helps fortify and repair k-fibers. This feature of Kif15 may help support robust k-fiber function and prevent chromosome segregation errors.

## Introduction

The mammalian mitotic spindle is a microtubule-based machine responsible for accurate chromosome segregation during cell division. In order for spindles to perform their essential function, they must be both robust and adaptable in a highly dynamic cellular environment. Kinetochore-fibers (k-fibers), microtubule bundles that link centrosomes to chromosomes, must therefore also be both robust and dynamic (Rieder, 1981; Hays *et al.*, 1982; McDonald *et al.*, 1992). Indeed, while single spindle microtubules turn over rapidly, k-fibers are more stable and turn over less frequently (Saxton *et al.*, 1984; Mitchison, 1989; McDonald *et al.*, 1992; Zhai *et al.*, 1995). Yet, k-fibers must not be too stable, or spindles may be unable to correct attachment errors, which can result in mis-segregated chromosomes (Lampson *et al.*, 2004). Regulation of microtubule lifetime is not the only way that spindles respond to competing demands: evidence also increasingly shows that the magnitude of force generation in the spindle balances robustness and dynamics in a myriad of ways (Pereira and Maiato, 2012; Elting *et al.*, 2014, 2017; Kajtez *et al.*, 2016; Long *et al.*, 2020; Suresh *et al.*, 2020). Yet, how k-fibers tune their mechanical integrity to maintain spindle structure and support accurate chromosome segregation remains unclear.

In mammalian mitotic spindles, kinetochore microtubules form a largely parallel array that spans from the kinetochore to the pole (Rieder, 1981; McDonald *et al.*, 1992). Thus, kinetochores hold together kinetochore microtubules at their plus-ends, while pole clustering forces, most notably provided by Nuclear Mitotic Apparatus protein (NuMA) and dynein, hold them together at their minus-ends (Merdes *et al.*, 1996; Gaglio *et al.*, 1997; Dionne *et al.*, 1999). Disrupting either kinetochores or poles alters spindle architecture as a whole, making it difficult to determine whether these end-based clustering forces are sufficient for stabilizing k-fibers. However, some evidence does suggest an important contribution of connections along k-fiber lengths to the mechanics of the k-fiber. K-fiber bundles appear to be much straighter and stiffer than individual microtubules (Goshima *et al.*, 2005), suggesting that the fibers somehow mechanically reinforce themselves. Indeed, when subjected to sufficient force the entire bundle often ruptures collectively (Long *et al.*, 2020). K-fiber reinforcement could come from connections between kinetochore microtubules themselves, or from associations with non-kinetochore microtubules. Electron microscopy, microneedle, and laser ablation experiments all indicate that nonkinetochore microtubules form mechanical connections with kinetochore microtubules within the k-fiber (Nicklas *et al.*, 1982; Hays and Salmon, 1990; McDonald *et al.*, 1992; Kajtez *et al.*, 2016; Elting *et al.*, 2017; Suresh *et al.*, 2020). While it is now clear that these non-kinetochore-microtubules contribute to local force dissipation across the spindle, they may also play a role in mechanical stabilization of the k-fiber itself.

Despite this evidence that they do so, it is not yet clear *how* interactions along the lengths of k-fiber microtubules, such as those through microtubule crosslinkers, contribute to the dynamic behavior and temporal longevity of the k-fiber as a whole (Elting *et al.*, 2018). A handful of microtubule-associated proteins (MAPs), such as the kinesin-12 Kif15 (Sturgill *et al.*, 2014) and its regulator TPX2 (Bird and Hyman, 2008; Sturgill *et al.*, 2014; Mann *et al.*, 2017), the clathrin/chTOG/TACC3 complex (Royle *et al.*, 2005; Booth *et al.*, 2011; Cheeseman *et al.*, 2013; Nixon *et al.*, 2015), HURP (Silljé *et al.*, 2006; Tsuchiya *et al.*, 2018), and kinesin Kif18A (Mayr *et al.*, 2007; Ye *et al.*, 2011) have been shown to localize preferentially to k-fibers and/or to stabilize them, thereby promoting efficient and reliable mitotic progression. However, their precise contributions to k-fiber mechanics and force production remain an open question.

The kinesin-12 Kif15 is well-positioned to shape k-fiber mechanics through both passive cross-linking and active force production. Like the kinesin-5 Eg5, a plus-end-directed motor responsible for separating poles in prometaphase to establish spindle bipolarity (Kapoor *et al.*, 2000), Kif15 is capable of crosslinking microtubule pairs and sliding antiparallel microtubules apart (Kapitein *et al.*, 2005; Drechsler *et al.*, 2014; Reinemann *et al.*, 2017). Through this mechanism, Kif15 can antagonize inward forces generated by minus-end-directed motors, thereby enforcing spindle bipolarity, and separating centrosomes in cells adapted to grow in the presence of Eg5 inhibitors (Raaijmakers *et al.*, 2012; Sturgill and Ohi, 2013). This redundancy between Kif15 and Eg5 is of interest as a potential factor in the disappointing ineffectiveness thus far of Eg5 inhibitors as cancer therapeutics (Tanenbaum *et al.*, 2009; Rath and Kozielski, 2012; Milic *et al.*, 2018; Dumas *et al.*, 2019). It is particularly notable since Kif15 overexpression results in lagging chromosomes (Malaby *et al.*, 2019), and is upregulated in a number of cancers (Ma *et al.*, 2014; Wang *et al.*, 2017; Qiao *et al.*, 2018; Yu *et al.*, 2019; Zhao *et al.*, 2019; Sun *et al.*, 2020; Terribas *et al.*, 2020). Under normal circumstances, however, Kif15 localizes predominantly to k-fiber microtubules. Its function on k-fibers is largely unexplored, although it is presumably from this location that the motor slides antiparallel microtubules apart.

Overall, the localization and known functions of Kif15 indicate that it might contribute to the mechanical properties of k-fibers. In addition, some evidence suggests that Kif15 coordinates the movement of neighbor sister kinetochores within the spindle (Vladimirou *et al.*, 2013). While it remains unclear whether Kif15 oligomerizes as a dimer or tetramer in vivo (Drechsler *et al.*, 2014; Mann *et al.*, 2017; Reinemann *et al.*, 2017), its second microtubule binding site, in addition to its motor domain, gives it inherent microtubule bundling ability even in a dimeric state (Sturgill *et al.*, 2014). Yet, Kif15 requires its binding partner, TPX2, for recruitment to the spindle (Tanenbaum *et al.*, 2009). Furthermore, *in vitro* data indicate that TPX2 increases Kif15 microtubule binding while decreasing motor activity (Drechsler *et al.*, 2014; Mann *et al.*, 2017; McHugh *et al.*, 2018), raising the question of how important motor activity is for its function on k-fibers.

To understand the dynamics and structural maintenance of the k-fiber, we mechanically disrupted k-fibers in the mitotic spindles of mammalian Ptk2 kidney epithelial cells. First, we used laser ablation to detach k-fibers from poles, disconnecting forces that may hold them together at their minus-ends. We found that the microtubules of these detached k-fibers often splay apart, and that this splaying results from minus-end clustering forces that can outcompete bundling forces. Next, to probe the role of Kif15 in k-fiber integrity, we tested the effect on k-fiber splaying of two Kif15 inhibitors with different mechanisms. Ablation experiments in cells treated with GW108X, which inhibits Kif15’s microtubule crosslinking activity, revealed a heightened sensitivity to these minus-end-directed forces. In contrast, we did not observe this effect in cells treated with Kif15-IN-1, which inhibits Kif15’s motor activity. The data presented here suggest that, independently of its motor activity, Kif15 mechanically reinforces k-fiber structure by binding and crosslinking k-fiber microtubules. The bundling forces provided at least in part by Kif15 are of a strength that allows them to compete with minus-end clustering forces, suggesting they play a mechanical role in preserving the integrity of k-fibers, thus supporting accurate chromosome alignment and segregation.

## Results and Discussion

### K-fibers are dynamically held together along their lengths, by connections that can be disrupted and reformed

In metaphase Ptk2 cells expressing GFP-α-tubulin, we performed laser ablations to remove forces holding k-fibers together at minus-ends and determine the degree to which k-fibers remain cohesive along their lengths (Figure 1A). The mechanical response to ablation commonly consisted of two readily observed behaviors, “poleward transport” and “splaying”. Poleward transport, a well-documented response to detached minus-ends (Elting *et al.*, 2014; Sikirzhytski *et al.*, 2014), followed nearly all ablations (Figure 1, B and C; Supplemental Figure S1F; Supplemental Videos SV1, SV2). We also frequently observe a behavior we term “splaying” in which the k-fiber stub’s microtubules dissociate along their lengths. While microtubule plus ends remain attached at the kinetochore, the minus ends of k-fiber stub microtubules come apart, losing the tightly-cohesive appearance typically associated with k-fibers (Figure 1A; Supplemental Video SV1). K-fiber stub splaying occurs following about half of all ablations (Figure 1D).

**Figure 1:**
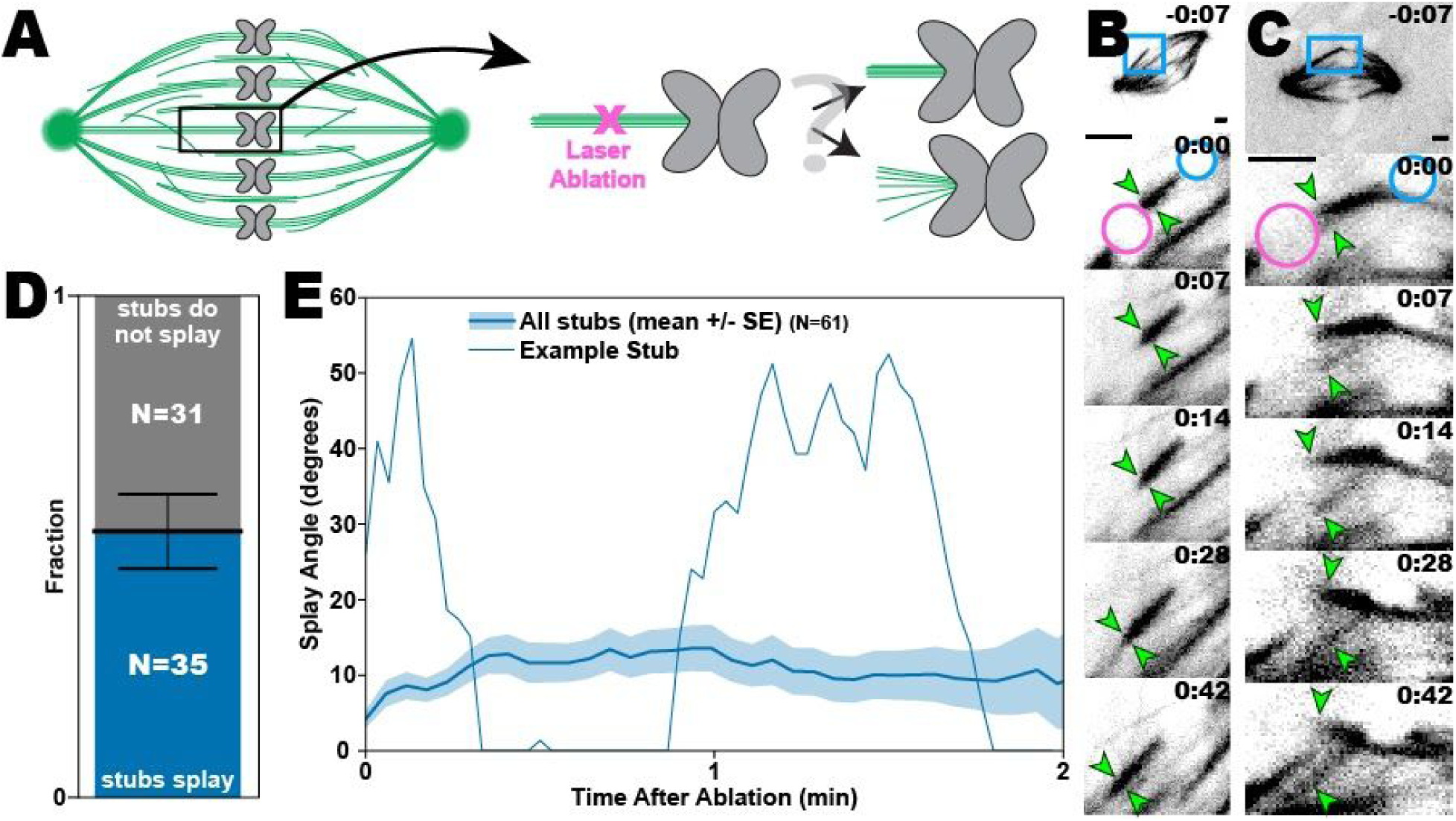
K-fiber microtubule bundles often splay apart following detachment from the pole by laser ablation. (A) Schematic of k-fiber ablation experiment. Following ablation (magenta ‘X’), k-fiber stubs demonstrate either “no splaying” (top) or “splaying” (bottom). (B&C) Typical examples of k-fiber stubs in Ptk2 cells, expressing GFP-α-tubulin, that do not splay (B) and that do splay (C). The ablation spot (magenta circle), k-fiber stub (green arrows), and chromosomes (blue circle) are labeled in the first post-ablation frame for both k-fiber stubs. All scale bars are 2 μm and timestamps are in min:sec. (D) Roughly half of ablated k-fiber stubs splay at some point during spindle repair. Error bar indicates the square root of the number of splayed events, as an estimate on the variation in this number assuming Poisson statistics. (E) Example trace (narrow blue line) showing the angle over time of a single ablated k-fiber stub, and the average (bold blue line) angle of all stubs. Although the average splay angle gradually increases to about 13°, before gradually decreasing, splaying varies widely among individual k-fiber stubs, with many splaying and “zipping up” multiple times and at different points during repair. Shaded region represents average +/- standard error on the mean.

When stubs splay, they typically separate into only two microtubule bundles (Figure 1C). The average amount of splaying increases for the first ∼20 s after ablation and remains at a steady average level after that (Figure 1E). However, the timing, duration, and repetition of these splaying events is highly variable among individual k-fiber stubs. For instance, after a k-fiber stub splays, the splayed microtubule bundles will usually “zip up”, reforming into a single tightly-bound k-fiber, a process that appears to occur simultaneously all along the k-fiber stub’s length. While this connection visually appears complete, it is often not final, as many k-fiber stubs undergo multiple splay-zip cycles during spindle repair (Figure 1E). Therefore, even though the collective average splay angle provides insight into the overall magnitude and timescale of k-fiber stub splaying, aggregate metrics obscure the significant temporal variability of behavior among individual k-fiber stubs (Figure 1E). This variation suggests an active competition between forces that bind neighboring k-fiber microtubules along their lengths and forces that pull on microtubule minus-ends.

### Minus-end clustering forces can overpower bundling forces to pry apart ablated k-fibers

Nearly all k-fiber stubs that splayed did so after the activation of machinery that pulls new minus-ends poleward (Figure 2A). Furthermore, the ability to maintain lateral inter-microtubule connections within k-fiber stubs is somewhat velocity-dependent, as k-fiber stubs that were splaying at any given moment experienced a greater poleward velocity than those that were not currently splaying (Figure 2B). These data suggest that forces that cluster minus-ends can also promote splaying, with increasing probability of splaying for k-fiber stubs experiencing stronger minus-end clustering forces.

**Figure 2:**
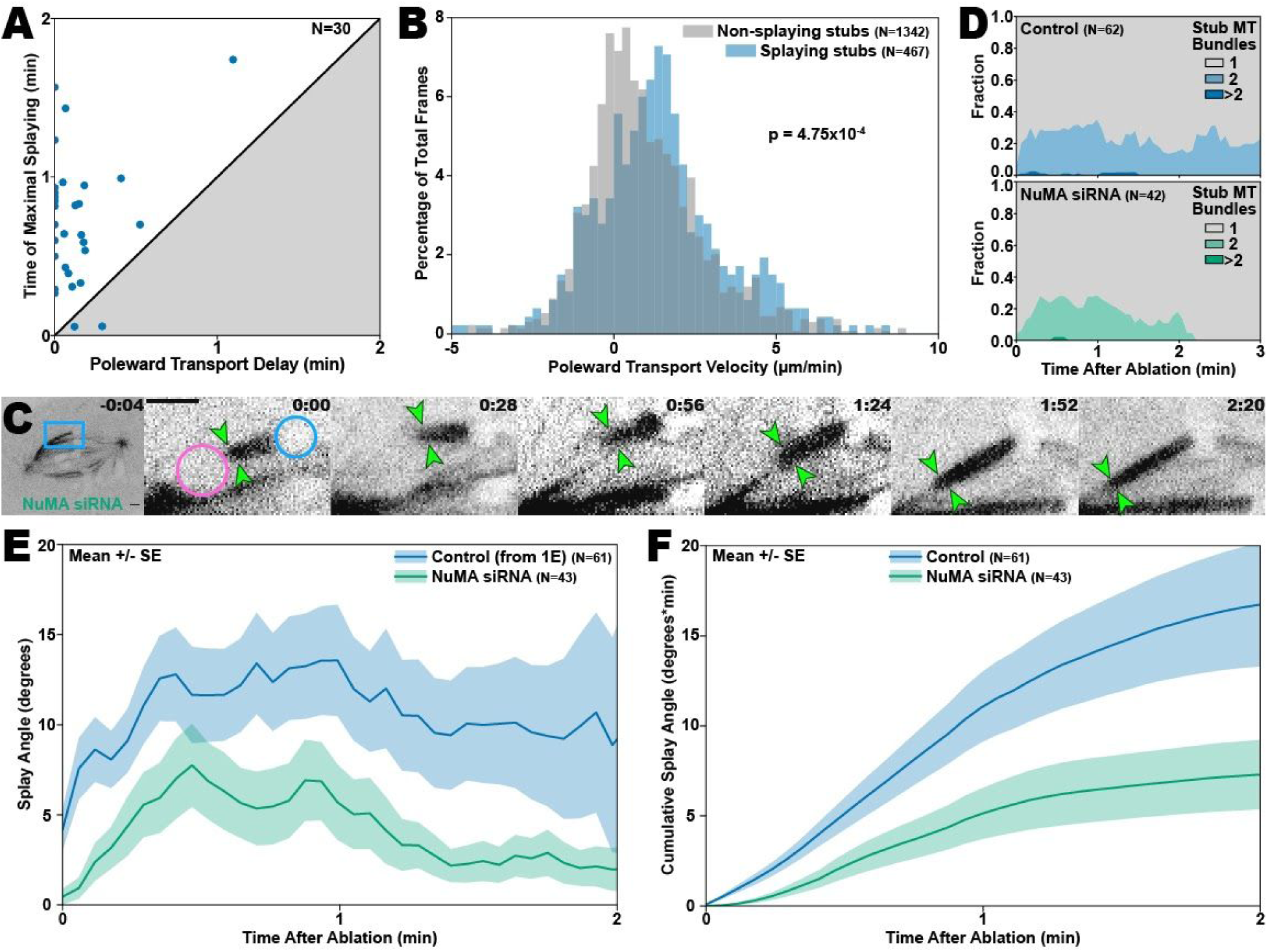
Minus-end clustering forces recruited in response to ablation increase the degree to which k-fiber bundles splay apart. (A) Maximal splaying typically occurs after the onset of poleward transport for untreated k-fiber stubs. Points represent individual k-fiber stubs. (B) K-fiber stubs undergoing rapid poleward transport are more likely to splay. Here the x-axis is the instantaneous poleward transport velocity and the y-axis is the percentage of time points in each histogram bin. (C) Example of an ablated k-fiber stub in a spindle of a Ptk2 cell expressing GFP-α-tubulin treated with NuMA siRNA. Following ablation, the k-fiber stub is transported poleward without splaying. The ablation spot (magenta circle), k-fiber stub (green arrows), and chromosomes (blue circle) are labeled in the first post-ablation frame. All scale bars are 2 μm and timestamps are in min:sec. (D) K-fiber stubs in spindles treated with NuMA siRNA (bottom, green) splay later than k-fiber stubs from control cells (top, blue) and do not stay splayed for as long. The y-axis represents the fraction of k-fiber stubs at each timepoint, consisting of 1, 2, or more than 2 visibly distinguishable microtubule bundles. (E) Splaying is reduced in NuMA siRNA treated cells compared with untreated cells. Shaded regions represent average +/- standard error on the mean. Note: these data for control k-fiber stubs were also shown in Fig. 1E, above. (F) Total cumulative splaying, measured as the integration of data shown in (E), is lower in cells treated with NuMA siRNA. Shaded regions represent average +/- standard error on the mean.

Nuclear mitotic apparatus protein (NuMA) has been shown to be required for rapid and robust poleward transport of loose spindle microtubule minus ends, ensuring the maintenance of spindle poles (Elting *et al.*, 2014; Sikirzhytski *et al.*, 2014). NuMA is recruited to these minus ends when they appear and in turn recruits cytoplasmic dynein and dynactin (Hueschen *et al.*, 2017), coalescing NuMA/dynein/dynactin complexes to the ablated k-fiber stub minus-end. Dynein motors, which are minus-end-directed, bind to another spindle microtubule and exert minus-end-directed force on the NuMA-bound microtubule, thereby transporting the k-fiber stub poleward as its cargo (Elting *et al.*, 2014; Sikirzhytski *et al.*, 2014). Because of the apparent dependence of k-fiber stub splaying on poleward transport, we next investigated the potential role of NuMA in splaying. To do so, we depleted NuMA via siRNA and then performed the same ablation experiments as before in these NuMA knockdown spindles (Figure 2C). As in previous work (Elting *et al.*, 2017), we chose spindles that retained bipolarity, indicative of only partial NuMA depletion, to increase the chance that any effects that we observed would be the direct result of NuMA perturbation rather than indirect effects from altering overall spindle architecture. As expected due to NuMA’s known involvement in poleward transport (Elting *et al.*, 2014; Hueschen *et al.*, 2017), poleward transport begins later and is more gradual for ablated k-fiber stubs in NuMA siRNA cells compared to those in control cells (Supplemental Figure S1D).

Based on NuMA’s established role as a mitotic spindle crosslinker, as well as its role anchoring k-fibers in the spindle body (Gaglio *et al.*, 1996; Elting *et al.*, 2017; Supplemental Figure S1A), we initially hypothesized that NuMA might itself mechanically bridge neighbor microtubules in the k-fiber. Strikingly, NuMA depletion resulted in more robust k-fiber bundling, as splaying of ablated k-fiber stubs is much more gradual initially and remains reduced throughout spindle repair (Figure 2D-F). Thus, it seems that the NuMA/dynein/dynactin complex, functioning as a minus-end-directed force generator, primarily works to pry ablated k-fiber stubs apart rather than hold them together. We hypothesize that this occurs when some of the multiple dynein motors recruited to the ablated minus-end bind to different track microtubules that are not precisely parallel. These data reinforce the hypothesis that minus-end-directed forces involved in poleward transport frequently contribute to longitudinal dissociation of microtubules within the k-fiber from each other.

### Kif15 binds and crosslinks k-fiber microtubules to promote k-fiber cohesion

We next used pharmacological inhibition to test for a potential role in k-fiber cohesion of Eg5 or Kif15, both plus-end-directed crosslinking motors. Based on the fact that poleward transport has been shown to be slightly faster but largely unchanged with Eg5 inhibition (Elting *et al.*, 2014), we expected that the splaying behaviors of ablated k-fiber stubs would remain unchanged (or potentially even increase) with the inhibition of Eg5. Indeed, both the poleward transport and splaying of ablated k-fiber stubs in STLC-treated spindles are generally similar to that of k-fiber stubs in untreated spindles (Supplemental Figure S1D-F). On the other hand, pharmacological inhibition of Kif15 revealed a more specific role for this motor in k-fiber mechanics. For these experiments, we used two Kif15 inhibitors that act by different mechanisms. Kif15-IN-1 arrests the motor in a microtubule-bound state, and thus blocks motility without disrupting its ability to crosslink microtubules (Milic *et al.*, 2018). By contrast, GW108X prevents the motor domain of Kif15 from binding the microtubule track, thus preventing the formation of microtubule crosslinks (Dumas *et al.*, 2019).

Unexpectedly due to Kif15’s plus-end-directed motor activity, the two Kif15 inhibitors, both of which cause loss of motor function, significantly perturb spindle repair. Poleward transport is more delayed than in control spindles with both inhibitors, and in the case of GW108X, is even more delayed than in NuMA siRNA spindles (Figure 3A and B; Supplemental Figure S1E). Under both Kif15 inhibition conditions, the spindle occasionally fails to repair altogether during observation, leaving the k-fiber still unattached to the pole at the end of filming (Supplemental Figure S1F). Although poleward transport of severed microtubule minus-ends has been strongly coupled with minus-end-directed force production by dynein (Elting *et al.*, 2014; Sikirzhytski *et al.*, 2014; Hueschen *et al.*, 2017), a few mechanisms might help explain the apparent additional requirement for Kif15, a plus-end-directed motor, in efficient poleward transport. For instance, Kif15 may act on anti-parallel overlaps near the middle of the spindle to help transport k-fiber stubs poleward following ablation. Alternatively, Kif15-mediated crosslinking might help mechanically anchor track microtubules on which the NuMA/dynein complex walks when transporting ablated k-fiber stubs. Finally, it is also possible that effective poleward transport of ablated k-fibers requires normal k-fiber architecture, which may be disrupted when Kif15 is inhibited (see below). Further investigation will be needed to discern the role of Kif15 in minus-end-directed force generation and the mechanism by which this force is produced.

**Figure 3:**
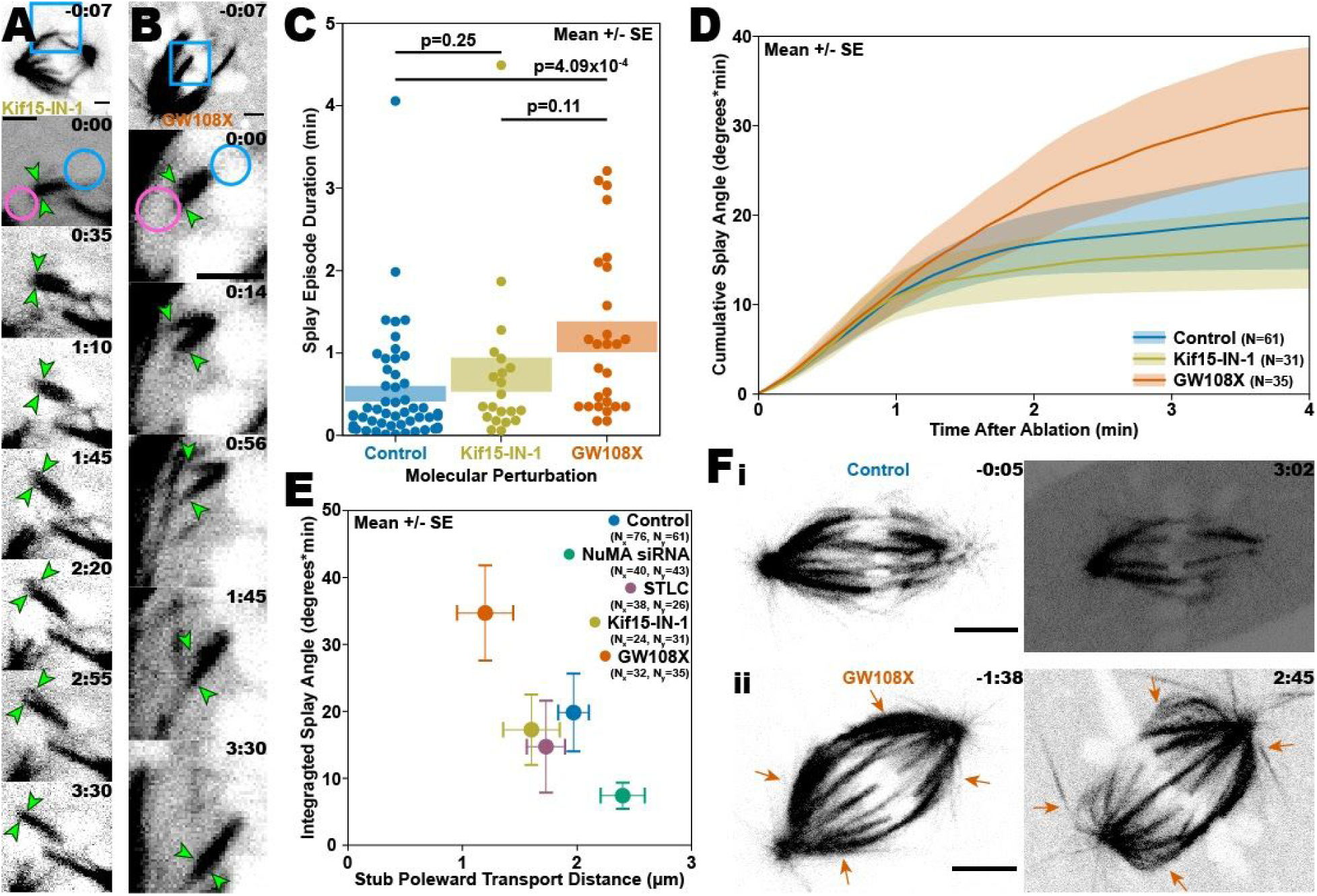
Crosslinking from Kif15 mediates the mechanical integrity of the k-fiber. (A & B) Typical examples of behavior of ablated k-fibers in spindles of Ptk2 cells treated with Kif15 inhibitors. The poleward transport of k-fiber stubs from Ptk2 cells treated with Kif15-IN-1 (A) is occasionally drastically delayed or entirely absent, and, in GW018X-treated cells (B), many of these k-fiber stubs that are not reincorporated into the spindle splay. The ablation spot (magenta circle), k-fiber stub (green arrows), and chromosomes (blue circle) are labeled in the first post-ablation frame of each panel. All scale bars are 2 μm and timestamps are in min:sec. (C) Kif15 inhibition by GW108X prolongs splaying. The y-axis is the length of time between when a k-fiber stub begins splaying and when the k-fiber stub is “zipped up”. Shaded regions represent average +/- standard error on the mean and numbers at the top of the plot represent p-values. (D) Total cumulative splaying, measured by integrating the k-fiber stub splay angle over time, is greater for k-fiber stubs in cells treated with GW108X (which inhibits crosslinking) but is about the same for k-fiber stubs in cells treated with KIF15-IN-1 (which solely inhibits motor activity), when compared to control k-fiber stubs. Shaded regions represent average +/- standard error on the mean. (E) Although treatment with GW108X reduces the poleward transport of ablated k-fiber stubs, relative to k-fiber stubs in untreated cells, these stubs still exhibit greater total splaying. In contrast, treatment with KIF15-IN-1 does not alter the total amount of splaying. Whiskers represent average +/- standard error on the mean. (F) Unablated k-fibers in control spindles usually remain tightly bound throughout metaphase in control spindles (i), while outer k-fibers (arrows) in spindles from GW108X-treated cells (ii) sometimes buckle and splay, while still attached at both the pole and kinetochore. Due to photobleaching, brightness and contrast of later time points is adjusted compared to earlier ones. Scale bars are 5 μm and timestamps are in min:sec.

Since Kif15 inhibition seems to slow and delay poleward transport of ablated k-fiber stubs, one would expect some reduction of splaying when compared with k-fiber stubs in control spindles, as we saw in the NuMA depletion experiments. Instead, stub splaying in cells treated with Kif15-IN-1 occurs with roughly the same frequency, duration, and magnitude as in untreated cells, while GW108X treatment exacerbates splaying (Figure 3, C-E; Supplemental Figure S1, C and F). For instance, when ablated k-fiber stubs in GW108X-treated cells splay, they often remain splayed for longer, a phenomenon far less pronounced in cells treated with Kif15-IN-1 (Figure 3C). On average, these k-fiber stubs also undergo more cumulative splaying over the course of the entire spindle repair process, demonstrating that Kif15 crosslinking is instrumental in the maintenance and reformation of inter-microtubule bonds within ablated k-fiber stubs (Figure 3D). The appearance of this effect specifically following exposure to GW108X, but not KIF15-IN-1, suggests that the relevant function of Kif15, as it relates to k-fiber stub splaying, is its inter-microtubule crosslinking. Furthermore, the increased total splaying for GW108X-treated k-fiber stubs occurs despite decreased poleward transport, as shown by tracking the kinetochore’s poleward displacement over the same period (Figure 3E). These data further point to the conclusion that the pronounced splaying results specifically from the loss of Kif15 crosslinking, independent of the effects of GW108X treatment on poleward transport. We see this dependence of Kif15 crosslinking even in unsevered k-fibers when GW108X-treated spindles sometimes shorten or collapse during imaging (Supplemental Figure S1D). When k-fibers occasionally buckle during collapse, it frequently induces splaying in the middle of these fibers, resulting in microtubule bundles that are apparently connected at both the centrosome and kinetochore, but are clearly unattached along much of their lengths (Figure 3Fii). While we sometimes see untreated spindles shorten somewhat over the course of imaging, their k-fibers do not typically splay when this occurs (Figure 3Fi). Stabilization of Kif15’s microtubule-bound state appears to prevent this buckling and splaying, as we observed this behavior less frequently in control spindles and never observed it in cells treated with Kif15-IN-1. These results are consistent with Kif15’s role in stabilizing kinetochore-microtubules during metaphase (Gayek and Ohi, 2014).

Altogether, these data suggest that, in addition to contributing to outward spindle-axis force generation, Kif15 promotes the mechanical cohesion of k-fibers along their lengths by crosslinking microtubules to form and maintain tightly bound bundles. By doing so, Kif15 helps the k-fiber both maintain its structural integrity under duress and, when its characteristic structure has been compromised, efficiently reform into a cohesive parallel microtubule array (Figure 4). This function of Kif15 in k-fiber structural maintenance might be particularly critical in other mammalian cell types in which k-fibers face additional mechanical challenges and constraints, such as crowding from higher numbers of chromosomes. It will also be interesting to investigate whether this function is further supported by molecular redundancy from other crosslinkers that localize to k-fibers, such as the clathrin/TACC3/ch-TOG complex and HURP. These or other molecules might contribute additional support to the mechanical redundancy Kif15 brings to the k-fiber, with microtubules reinforcing each other to help ensure accurate spindle attachment and chromosome segregation. Such a balance between molecular force generators might help k-fibers to reshape themselves in response to spindle dynamics.

**Figure 4:**
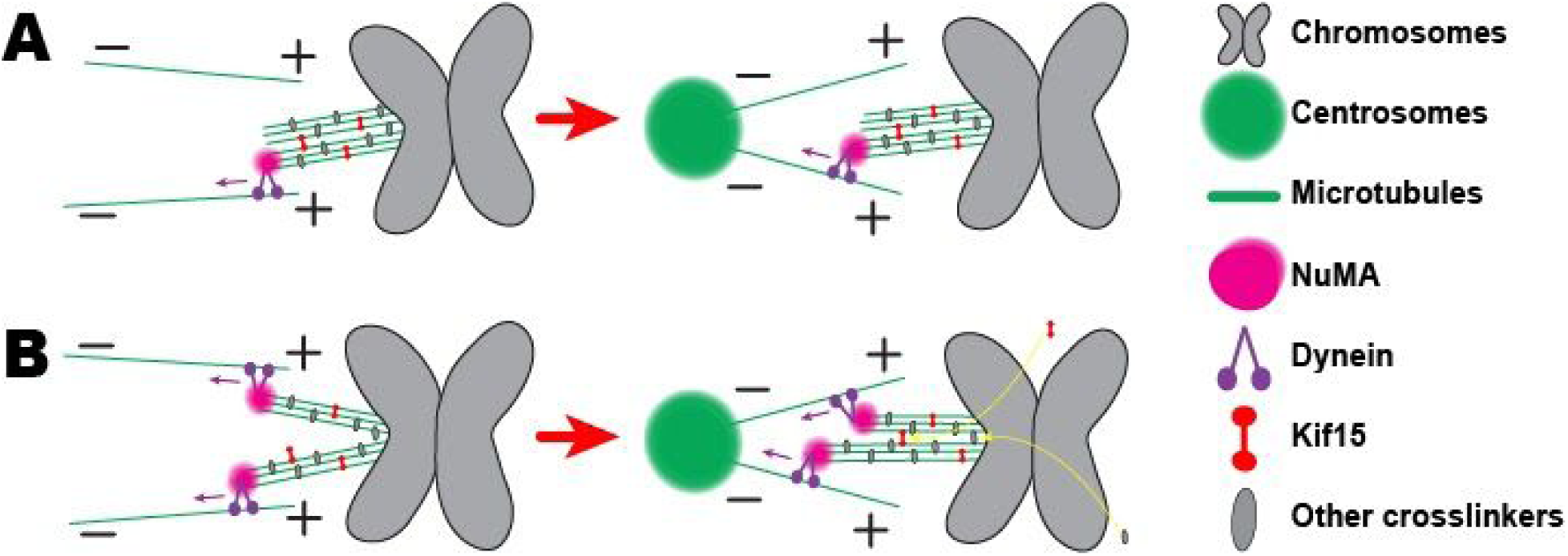
Kif15 crosslinking encourages the formation and maintenance of k-fiber microtubule cohesion. (A & B) During spindle repair, the microtubules of some k-fiber stubs remain tightly bound together, while others splay apart. (A) Microtubule pairs crosslinked by Kif15 often withstand spindle forces during poleward transport. (B) When k-fiber stubs splay, Kif15 re-crosslinks microtubules to “zip up” splayed k-fiber stubs. When Kif15 microtubule binding is perturbed, k-fiber stubs are more likely to splay even before, or in the absence of, poleward transport.

## Materials & Methods

### Cell culture

Wild type Ptk2 cells and Ptk2 stably expressing GFP-α-tubulin (both a gift of Sophie Dumont, UCSF) were cultured in MEM (Genesee 25-504 or Fisher 11095080) supplemented with non-essential amino acids (Genesee 25-536), sodium pyruvate (Genesee 25-537), penicillin/streptomycin (Genesee 25-512), and heat-inactivated fetal bovine serum (Genesee 25-514H). For live imaging, cells were plated in 9.6 cm^2^ glass-bottom dishes treated with poly-D-lysine (MatTek P35GC-1.5-14-C) in 2 mL of MEM-complete (described above). For immunofluorescence experiments, cells were plated on #1.5 25 mm coverslips (HCl cleaned and treated with poly-L-lysine) in a 6-well plate.

### Drug treatment

For treatment with KIF15-IN-1 (Apex Bio), inhibitor was added to plated cells at a final concentration 12.5 μM (from 10mM stock in DMSO) and imaging was performed 1-2 hours after treatment. For treatment with GW108X (custom synthesis as previously described (Dumas *et al.*, 2019)), inhibitor was added to plated cells at a final concentration 25 μM (from 10mM stock in DMSO) and imaging was performed within 1 hour of treatment. For treatment with S-trityl-L-cysteine (STLC) (Sigma-Aldrich 164739-5G), inhibitor was added to plated cells at a final concentration 5 μM (from 20mM stock in DMSO) and imaging was performed within 1 hour of treatment. Using these conditions, morphological signs of both KIF15-inhibition and Eg5-inhibition in spindles (such as shortened spindles) were detectable (Fig. 1C), but many spindles still displayed fairly normal overall bipolar morphology. Although monopoles and fully-collapsed bipoles were often observed, further verifying successful treatment, we did not perform ablation experiments on these cells.

### Transfections & siRNA

For NuMA siRNA experiments, GFP-α-tubulin Ptk2 cells were transfected with siRNA for NuMA as previously described (Udy *et al.*, 2015; Elting *et al.*, 2017). The sequence for our siNuMA was 5’-GCATAAAGCGGAGACUAAA-3’, designed based on the Ptk2 transcriptome, and Oligofectamine (Invitrogen 12252-011) was used for transfection reagent (Udy *et al.*, 2015; Elting *et al.*, 2017). Transfected plates were either fixed or imaged live 48-96 hr following treatment, with most experiments conducted on the third day after transfection. We verified our knockdown by observing NuMA expression and localization in fixed cells via immunofluorescence assay (Fig. 1A). In live cells, we also frequently observed spindle abnormalities, such as splayed or multiple poles, further verifying successful NuMA knockdown. However, we chose cells with bipolar spindles for ablation experiments, in order to focus on a direct effect of NuMA rather than effects of perturbing overall spindle architecture.

### Immunofluorescence

Immunofluorescence was used to verify knockdown siRNA of NuMA. For fixation, cells were treated for three minutes with a solution of 95% methanol and 4.8mM EGTA. The following antibodies were used for these experiments: mouse anti-α-tubulin DM1α (1:500, Invitrogen 62204), rabbit anti-NuMA (1:400, Novus Biologicals NB500-174SS), human anti-centromere protein (CREST; 1:25, Antibodies Inc 15-234), fluorescent secondary antibodies (1:500, Invitrogen), and Hoechst 33342 (Invitrogen H3570). Coverslips were mounted on slides using Prolong Gold. Following fixation, cells were imaged using the confocal fluorescence microscope described below. For verifying knockdown, identical conditions (for fixation, staining, and imaging) were used to compare knockdown and control cells.

### Live cell imaging and laser ablation

For live cell imaging, cells were plated in 9.6 cm^2^ glass-bottom poly-D-lysine-coated dishes (MatTek P35GC-1.5-14-C) in 2mL of MEM-complete (described above) and left in an incubator at 37°C and 5% CO_2_ until time for imaging.

Live imaging and ablation experiments were performed essentially as previously described (Elting *et al.*, 2017). In this case, live imaging was performed on a Nikon Ti-E stand on an Andor Dragonfly spinning disk confocal fluorescence microscope; spinning disk dichroic Chroma ZT405/488/561/640rpc; 488 nm (50 mW) diode laser (240 ms exposures) with Borealis attachment (Andor); emission filter Chroma Chroma ET525/50m; and an Andor iXon3 camera. Imaging was performed with a 100x 1.45 Ph3 Nikon objective and a 1.5x magnifier (built-in to the Dragonfly system). Frames were collected every 0.3-4.0 s for up to ∼4 minutes after ablation. Targeted laser ablation was performed using an Andor Micropoint attachment with galvo-controlled steering to deliver 20-30 3 ns pulses at 20 Hz of 551 nm light. Andor Fusion software was used to control acquisition and Andor IQ software was used to simultaneously control the laser ablation system. For all experiments involving live cells, imaging was conducted on a closed stagetop incubator (Okolab), which maintains conditions of 30°C, 5% CO_2_ and humidity.

### Data analysis

FIJI was used to prepare all videos for analysis. This process consisted of cropping frames, adjusting brightness and contrast, and converting file types. We cropped GFP-α-tubulin ablation images, in order to more clearly track the positions and splay angles of ablated k-fiber stubs. Videos were saved as both a TIFF stack and an AVI copy for each. The ‘No Compression’ option in FIJI was used, when saving videos as AVI files. Linear adjustments were made to the brightness and contrast of immunofluorescence, cotransfection, and GFP-α-tubulin ablation videos. The brightness and contrast of all videos in each immunofluorescence and cotransfection data were scaled the same as all other videos in the set.

In videos of GFP-α-tubulin spindles, spindle length was measured using the ‘line’ tool in FIJI. These measurements were conducted at the beginning of imaging of the spindle, to prevent potential effects on spindle length of long term imaging. Poles were identified as the center of high-intensity circles at the spindle ends. The focus of radial microtubule bundles was designated as the pole position for poles that were out of focus or dim.

Following ablations, k-fiber stubs first drifted away from the pole, toward the spindle midzone, before being transported poleward. When a k-fiber stub was not visibly transported poleward during at least four minutes of imaging, this was counted as an instance in which spindle repair (and poleward transport) does not transpire. We define poleward transport onset as the time at which the distance between the ablated k-fiber stub’s kinetochore and pole is maximal. This was done via a tracking program, written in Python, for videos in which both the kinetochore and pole remain visible throughout. Poles were identified in a manner similar to that used to measure spindle length and kinetochore positions were defined as k-fiber stub plus-ends. Video playback was used, as many times as needed, to verify all pole and kinetochore position measurements.

Another tracking program, home written in Python, was used to measure ablated k-fiber stub splay angle. For videos in which the k-fiber stub was visible throughout, the splay angle was defined as the angle formed by the k-fiber stub’s plus-end and the minus-ends of the two most separated splayed microtubule bundles. Occasionally, microtubule bundles were not approximately straight. In this case, instead of clicking on the bundle’s minus-end, clicks were made somewhere along the bundle’s length, so that the measured splay angle more accurately reflects the angle between the bundles proximal to the kinetochore. All splay angle measurements are verified by video playback, as many times as needed. Splay durations were measured as the length of uninterrupted time for which a k-fiber stub was splayed at an angle greater than 15°. Before applying this splay angle threshold, splay angle vs time data were smoothed using a 7-point binomial filter (Marchand and Marmet, 1983; Aubury and Luk, 1996); Figure 2, A and B; Figure 3C; Supplemental Figure S1E), making it easier to identify the initiation and conclusion of each splaying event. This same filter was applied to the k-fiber stub displacement vs. time data for calculating the instantaneous velocity (Figure 2, A and B).

## Acknowledgments

We thank Sophie Dumont, Dick McIntosh, Christina Hueschen, Jon Kuhn, Alex Long, and Elting Lab members for helpful discussions about the work. We thank Sophie Dumont (UCSF) for the gift of Ptk2 cells expressing GFP-α-tubulin. We thank Eva Johannes and Mariusz Zareba of the North Carolina State University Cellular and Molecular Imaging Facility for helpful discussions and assistance with microscopy. MWE and MAB were supported by a Ralph E. Powe Junior Faculty Enhancement Award from the Oak Ridge Associated Universities. RO and ALS are supported by R01 GM086610. The authors declare no competing financial interests.

## Supplemental Information

### Supplemental Figures

**Supplemental Figure S1:**
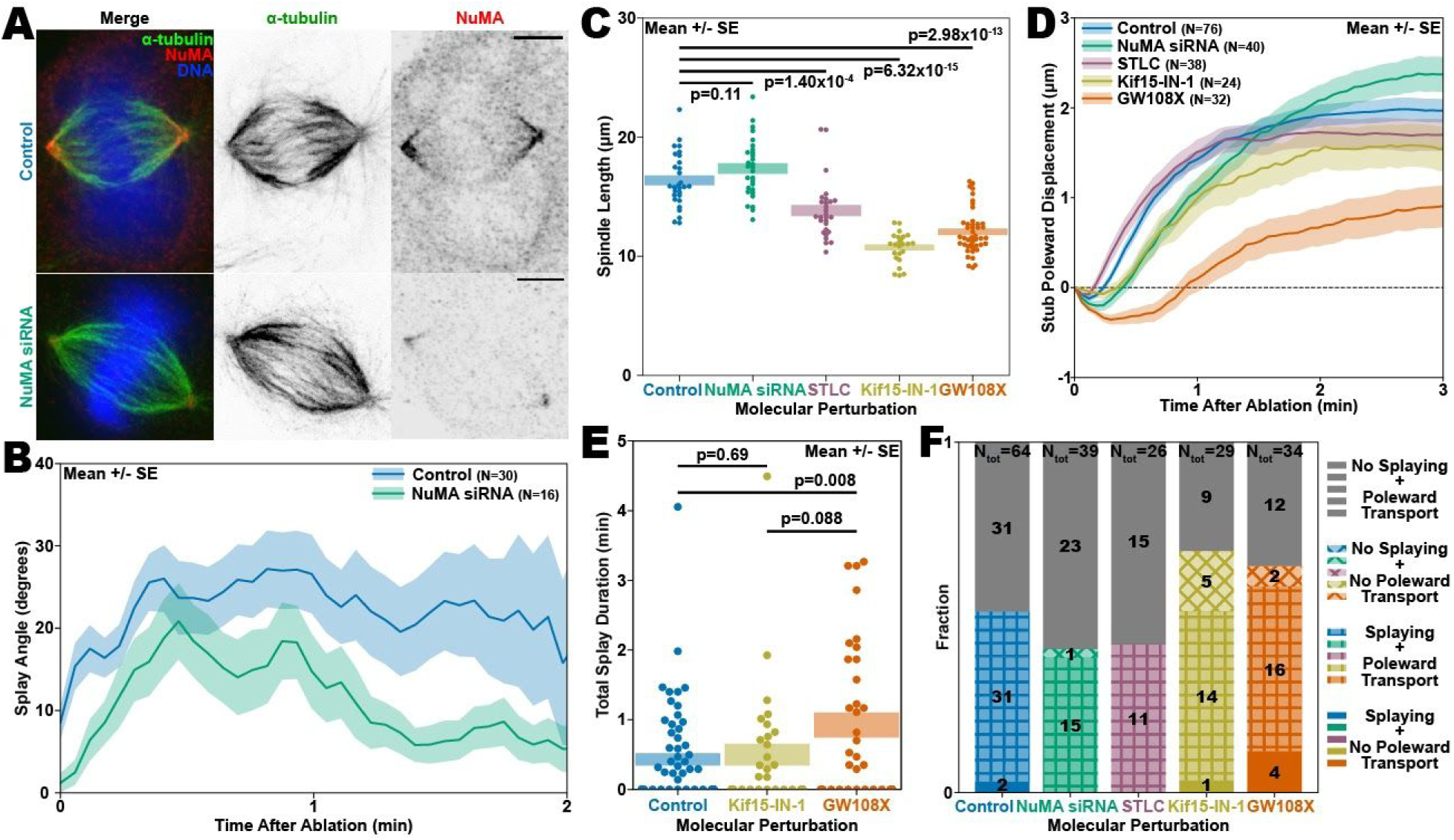
(A) In immunofluorescence experiments, NuMA siRNA reduces NuMA k-fiber localization, particularly near the poles. All scalebars are 5 μm. While this example shows a bipolar spindle, typical of those that we use for ablation experiments, we also saw many spindles with splayed poles or multipolar spindles when we treated with siRNA for NuMA. (B) NuMA siRNA reduces splaying in splaying k-fiber stubs, compared with splaying k-fiber stubs in untreated cells. In contrast with Fig. 2E, this panel includes only splaying k-fiber stubs, and demonstrates that the effects in Fig. 2E are from a lower proportion of k-fiber stubs that splay combined with less splaying in those that do splay. Shaded regions represent average +/- standard error on the mean. (C) Eg5 and Kif15 inhibition results in shortened spindles. Shaded regions represent average +/- standard error on the mean and numbers at the top of the plot represent p-values. (D) NuMA siRNA and Kif15-IN-1 inhibition cause modest delays in poleward transport of ablated k-fiber stubs, while inhibiting Kif15 with GW108X results in poleward transport with severe delays and decreased overall magnitudes. Shaded regions represent average +/- standard error on the mean. (E) Kif15 inhibition with GW108X increases the time that k-fiber stubs remain splayed during repair. Shaded regions represent average +/- standard error on the mean and numbers at the top of the plot represent p-values. (F) Categorization of k-fiber stubs based on overall behavior. Numbers represent the number of k-fiber stubs exhibiting each behavior for each molecular condition and the total number of observed k-fiber stubs in each condition is given at the top of the corresponding bars. Note the number of Kif15-inhibited k-fiber stubs that do not undergo poleward transport (diagonal cross hatching and solid colors, yellow and orange), as well as those in GW108X-treated cells that are not reincorporated, but nonetheless splay (solid orange).

### Supplemental Video Legends

**Supplemental Video SV1.** Typical example of an ablated k-fiber stub of a Ptk2 cell expressing GFP-α-tubulin which splays during poleward transport.

**Supplemental Video SV2.** Typical example of an ablated k-fiber stub of a Ptk2 cell expressing GFP-α-tubulin which does not splay during poleward transport.

**Supplemental Video SV3.** Typical example of an ablated spindle from a Ptk2 cell expressing GFP-α-tubulin and treated with GW108X. The ablated k-fiber stub splays before poleward transport, a behavior that is typical when treated with GW108X but very unusual in either cells treated with KIF15-IN-1 or in untreated control spindles.

